## Supplementary figures and images for "Molecular crowding in single eukaryotic cells: using cell environment biosensing and single-molecule optical microscopy to probe dependence on extracellular ionic strength, local glucose conditions, and sensor copy number"

### Supplementary Movie 1

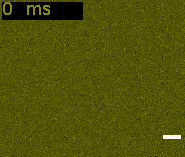
